# Coral: a web-based visual analysis tool for creating and characterizing cohorts

**DOI:** 10.1101/2021.05.26.445802

**Authors:** Patrick Adelberger, Klaus Eckelt, Markus J. Bauer, Marc Streit, Christian Haslinger, Thomas Zichner

## Abstract

**Summary:** A main task in computational cancer analysis is the identification of patient subgroups (i.e., cohorts) based on metadata attributes (patient stratification) or genomic markers of response (biomarkers). Coral is a web-based cohort analysis tool that is designed to support this task: Users can interactively create and refine cohorts, which can then be compared, characterized, and inspected down to the level of single items. Coral visualizes the evolution of cohorts and also provides intuitive access to prevalence information. Furthermore, findings can be stored, shared, and reproduced via the integrated session management. Coral is pre-loaded with data from over 128,000 samples from the AACR Project GENIE, The Cancer Genome Atlas, and the Cell Line Encyclopedia.

**Availability and Implementation:** Coral is publicly available at https://coral.caleydoapp.org. The source code is released at https://github.com/Caleydo/coral.

**Supplementary information:** Supplementary data are available at *Bioinformatics* online.

## 1 Introduction

A frequent goal in cancer research is the characterization of patient or sample groups based on a rich collection of experimental and metadata. A common approach is to define multiple cohorts (i.e., groups of items that share one or more characteristics), for instance, by splitting the samples based on their response to a treatment. One then aims to identify shared properties, find differences, and explore cohort characteristics in detail. Interactive visualization tools can assist users in creating, refining, and comparing cohorts based on attributes from high-dimensional space. Current tools, such as cBioPortal (Cerami *et al.*, 2012; Gao *et al.*, 2013), Composer (Rogers *et al.*, 2019), and CAVA (Zhang *et al.*, 2015), however, do not allow comparison or refinement of more than two cohorts, compare attributes only individually, or do not show the relationships between the cohorts created (for details see Supplementary Notes).

To address this deficiency, we present *Coral*, a web-based tool for interactively creating and characterizing multiple cohorts based on a large number of attributes (Fig. 1). Coral allows users to keep track of the cohort definition and refinement process (*cohort evolution*) by showing which attributes have been used to create cohorts and how they are related to each other. Additionally, analysts can use tailored visualizations to characterize the cohorts and explore them in detail. The integrated session management allows users to reproduce and share the cohorts they created during their analysis sessions (see Supplementary Notes).

**Fig. 1:**
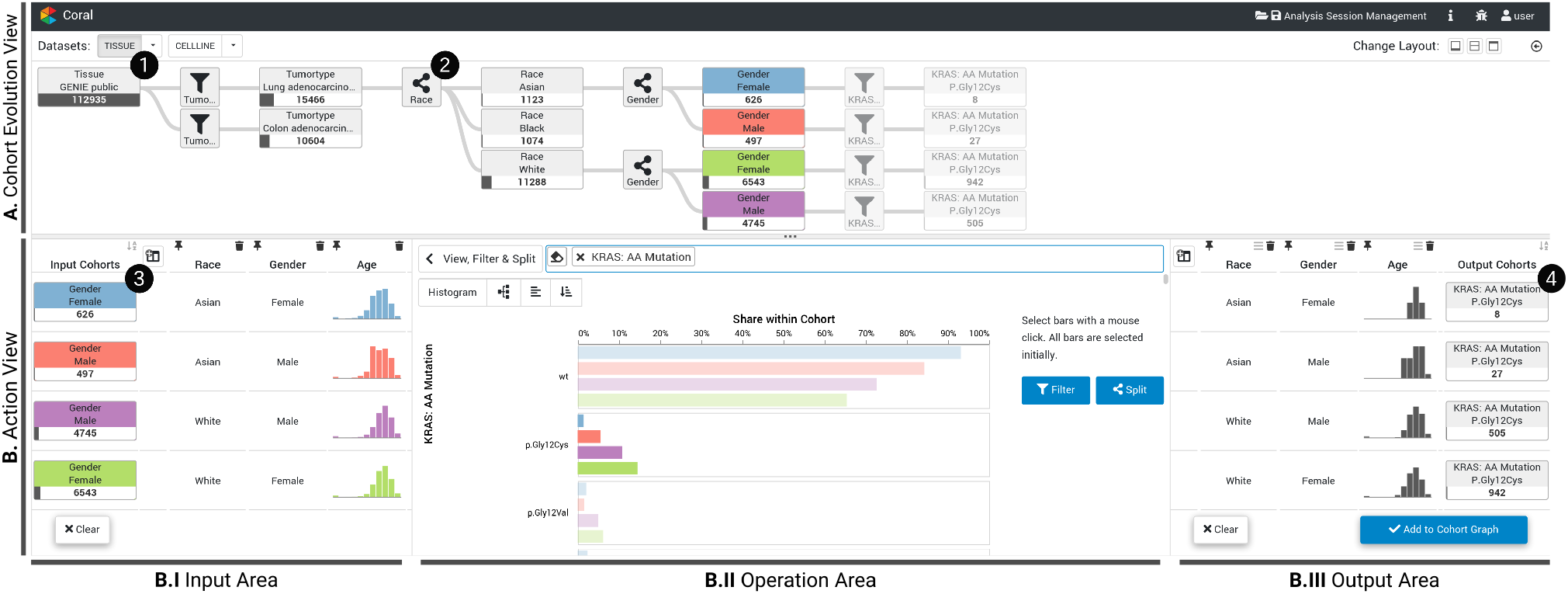
Using Coral to reproduce *KRAS^G12C^* mutation findings by Nassar *et al.* (2021). The *Cohort Evolution View* (A) shows that the user starts by filtering the AACR Project GENIE data to include only patients with either non-small cell lung cancer (NSCLC) or colorectal cancer. The user then splits the NSCLC cohort by race, and subsequently the resulting cohorts *Asian* and *White* by gender. The resulting cohorts are selected and highlighted (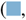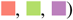) across all views. The *Input Area* (B.I) shows the cohorts’ distributions for race, gender, and age. Using *View, Filter & Split* in the *Operation Area* (B.II), the user subsequently assesses the four cohorts with respect to their frequencies of various *KRAS* mutations and selects the samples that contain *KRAS^G12C^*. By applying the *Filter* operation, the user generates the corresponding four output cohorts displayed in the *Output Area* (B.III).

We demonstrate the utility of our tool by reproducing findings from a recently published article about *KRASG12C* somatic mutations (Nassar *et al.*, 2021) and using a case study exploring different *KRAS* mutations across multiple tumor types (see Supplementary Notes, Figs. S3–S17, and Video).

## 2 Software Description

Coral is a client-server web-application that utilizes a Postgres database and is implemented in TypeScript and Python. Coral’s database contains mutation data from the AACR Project GENIE (AACR Project GENIE Consortium, 2017), mRNA expression, DNA copy number, and mutation data from The Cancer Genome Atlas (TCGA; https://cancergenome.nih.gov) and the Cell Line Encyclopedia (CCLE) (Barretina *et al.*, 2012), and two depletion screen datasets. In total, Coral’s database contains data of over 128,000 samples. Data pre-processing steps are described in the Supplementary Notes. The Coral interface follows an overview+detail approach in which two components are shown in a horizontal split-screen setup.

The **Cohort Evolution View** (upper panel) presents how all cohorts were generated as well as their relationships as a graph. Operations and cohorts are encoded as nodes connected by edges to represent the analysis flow (see Fig. 1A). The first cohort includes all items of the loaded dataset and is created automatically (Fig. 1 ❶). When the user selects a cohort in the graph, it is assigned a color that is used consistently in all visualizations. Selected cohorts are loaded into the **Action View** (lower panel), which allows users to perform cohort characterization and creation operations. New cohorts created by these operations are also added to the graph, which results in an iterative cohort definition and analysis workflow (see Supplementary Notes, Fig. S1). The Action View is divided into three areas: the *Input Area*, the *Operation Area*, and the optional *Output Area*, which is shown if the operation results in new cohorts (Fig. 1B). The *Operation Area* provides the following operations that can be applied to the input cohorts.

### View

In addition to the visualizations in the *Input Area* (Fig. 1B.I), which provide context to the input cohorts, the *View* operation is the main route to exploring the dataset and investigating how the values of one or more attributes are distributed across cohorts. The visualization is based on the number and type of attributes, as explained in Supplementary Table S1. **Filter & Split.** The *Filter & Split* operation is used to create cohorts from the loaded dataset. Fig. 1B shows four input cohorts being filtered using the categorical attribute *KRAS mutation status*. Users can adjust the selection, as illustrated in Fig. 1B.II, where, for each input cohort, the user selected all samples with a *KRASG12C* mutation. In contrast, the *Split* operation can be used to divide a cohort into multiple sub-cohorts, as shown in Fig. 1 ❷, where the NSCLC cohort is split based on *Race*.

### Compare

Additionally to comparing cohorts using the visualizations in the *Input Area* and the *View* operation, Coral offers a *Compare* operation to test for statistically significant differences. Supplementary Figs. S6– S7 show that the NSCLC and the colorectal cancer cohorts in Fig. 1 differ significantly in their age and gender distributions. The compare operation integrates the TourDino view (Eckelt *et al.*, 2019) to confirm visual patterns. Cohorts are tested pairwise for differences. The test results are summarized in a matrix showing each p-value. The applied test, its score, and a visualization are shown on demand.

### Prevalence

Prevalence is the proportion of items with a certain characteristic in a cohort. In Fig. 1, the proportion of patients with a *KRAS^G12C^* mutation (*sample cohort* ❹) among female Asian patients with NSCLC (*reference cohort* ❸) is 8*/*626, which equals a prevalence of about 1.3%. Coral provides a dedicated analysis view to assess prevalence estimates (see Supplementary Fig. S13). After selecting the *sample cohort* with items that have the characteristic of interest, the user can flexibly define the *reference cohort* by applying or skipping *Filter & Split* operations used to create the *sample cohort*. The cohorts’ sizes and the resulting prevalences are then displayed in a bar chart.

### Inspect Items

To identify outliers or to assess single data points, users need to move from the aggregated level down to the level of individual items. We use the tabular visualization technique Taggle (Furmanova *et al.*, 2020) for visualizing the items of the cohorts. Users can select attributes, whose data is displayed in the table, and sort, filter, and group the data (see Supplementary Fig. S16).

## 3 Conclusion and Future Work

Coral is an analysis tool for interactively creating cohorts based on a range of attributes, and visually represents the evolution of cohorts in an overview graph. The cohorts can be characterized, either by size or by a rich set of experimental and metadata. Users are able to gain insights into the cohort prevalences and also to drill down to the level of single-item information. In the future, we plan to further improve Coral by, for instance, supporting longitudinal data that contains multiple samples per patient from different time points. Furthermore, the current focus of Coral is on cancer genomics, however, technically and conceptually, Coral can be applied to other fields. We plan to support the upload of custom data and connections to additional databases in the future. By doing so, Coral will be able to cater the needs of an even broader range of users. For more details on future work see Supplementary Notes.

## Acknowledgements

We thank Andreas Wernitznig, Alex Lex, and the datavisyn GmbH team, in particular Holger Stitz, for their feedback and implementation support. The authors acknowledge the American Association for Cancer Research and its financial and material support in the development of the AACR Project GENIE registry, and members of the consortium for their commitment to open data. Interpretations are the responsibility of study authors.

## References

AACR Project GENIE Consortium (2017). AACR Project GENIE: powering precision medicine through an international consortium. Cancer discovery, 7(8), 818–831.

Barretina, J. et al. (2012). The Cancer Cell Line Encyclopedia enables predictive modelling of anticancer drug sensitivity. Nature, 483(7391), 603–607.

Cerami, E. et al. (2012). The cBio Cancer Genomics Portal: An Open Platform for Exploring Multidimensional Cancer Genomics Data. Cancer Discovery, 2(5), 401–404.

Eckelt, K. et al. (2019). TourDino: A Support View for Confirming Patterns in Tabular Data. In Proceedings of the EuroVis Workshop on Visual Analytics, pages 7–11.

Furmanova, K. et al. (2020). Taggle: Combining overview and details in tabular data visualizations. Information Visualization, 19(2), 114–136.

Gao, J. et al. (2013). Integrative Analysis of Complex Cancer Genomics and Clinical Profiles Using the cBioPortal. Science Signaling, 6(269), pl1

Nassar, A. H. et al. (2021). Distribution of KRASG12C Somatic Mutations across Race, Sex, and Cancer Type. New England Journal of Medicine, 384(2), 185–187.

Rogers, J. et al. (2019). Composer — Visual Cohort Analysis of Patient Outcomes. Applied Clinical Informatics, 10(02), 278–285.

Zhang, Z. et al. (2015). Iterative cohort analysis and exploration. Information Visualization, 14, 289–307.

