## Supplementary Notes for "Coral: a web-based visual analysis tool for creating and characterizing cohorts"

**Supplementary References** **20**

### 1 Supplementary Figures

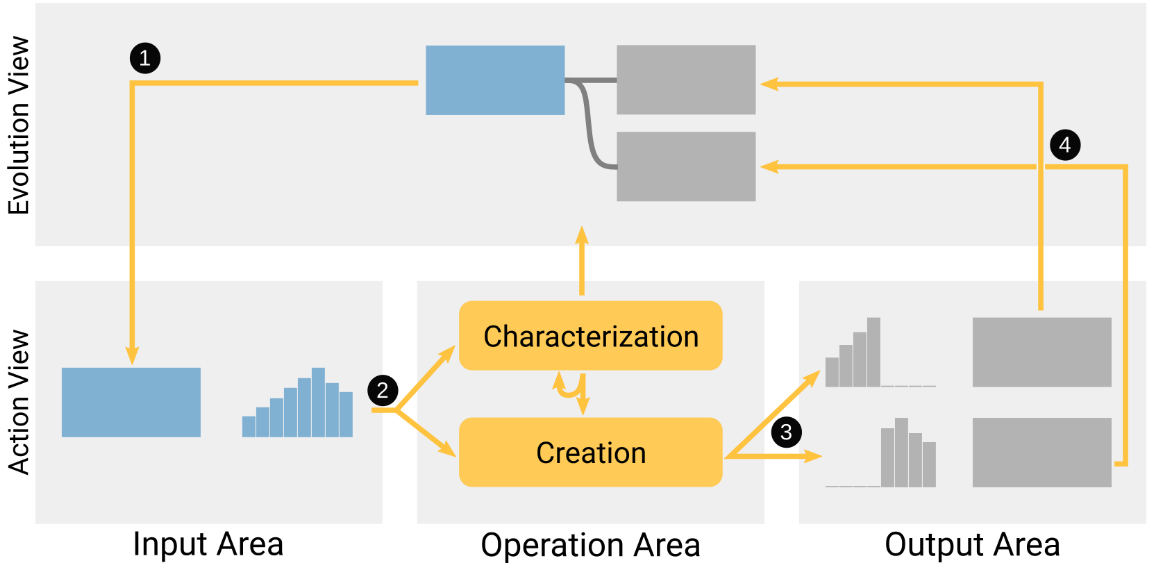

**Supplementary Figure S1**: **Coral workflow.** Cohort creation workflow in Coral. The selected, blue cohort (■) is added to the *Input Area* (1) and split in the *Operation Area* (2). The two resulting subsets define two output cohorts (3) that are added to the *Cohort Evolution View* (4).

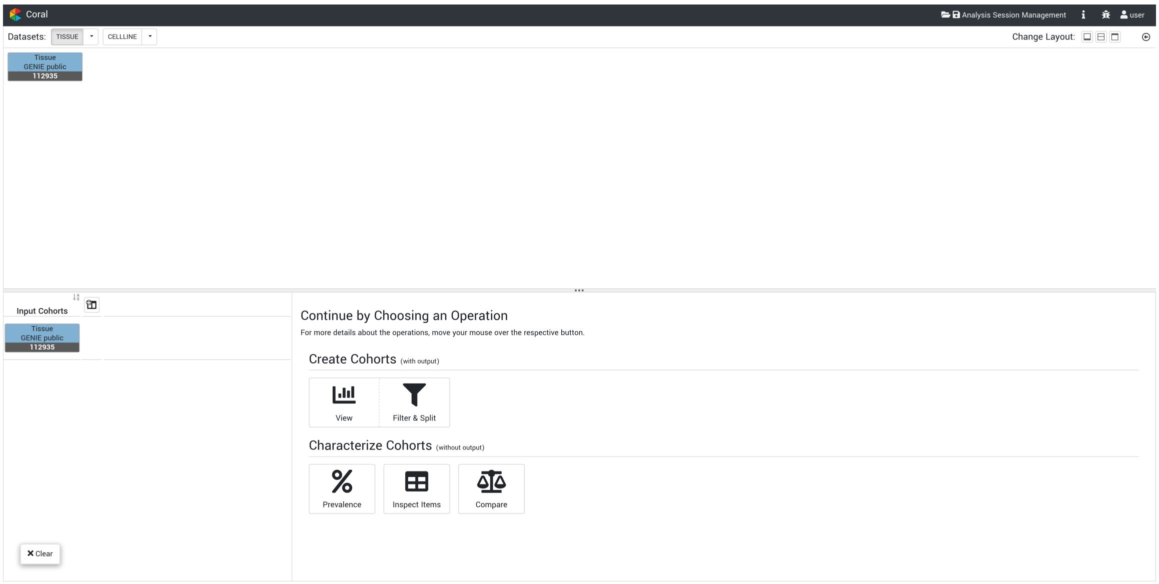

**Supplementary Figure S2**: **Operation selection.** The *Operation Area* allows selecting which operation to perform. The operations available are grouped by type. Available are: the *View* operation with integrated *Filter & Split* operations for cohort creation and the *Prevalence*, *Inspect Items*, and *Compare* operations for characterization of cohorts.

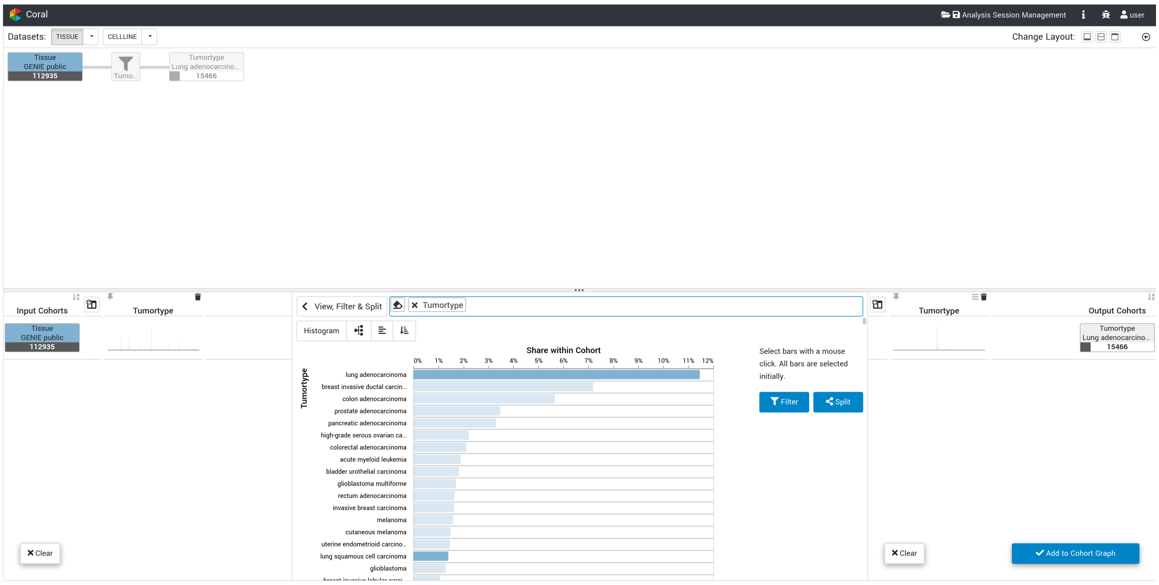

**Supplementary Figure S3**: **Case study 1:** Filtering of the starting cohort (■), which includes all AACR Project GENIE samples, down to non-small cell lung cancer (NSCLC) samples. Included tumor types are: *lung adenocarcinoma*, *lung squamous cell carcinoma*, and *non-small cell lung cancer*.

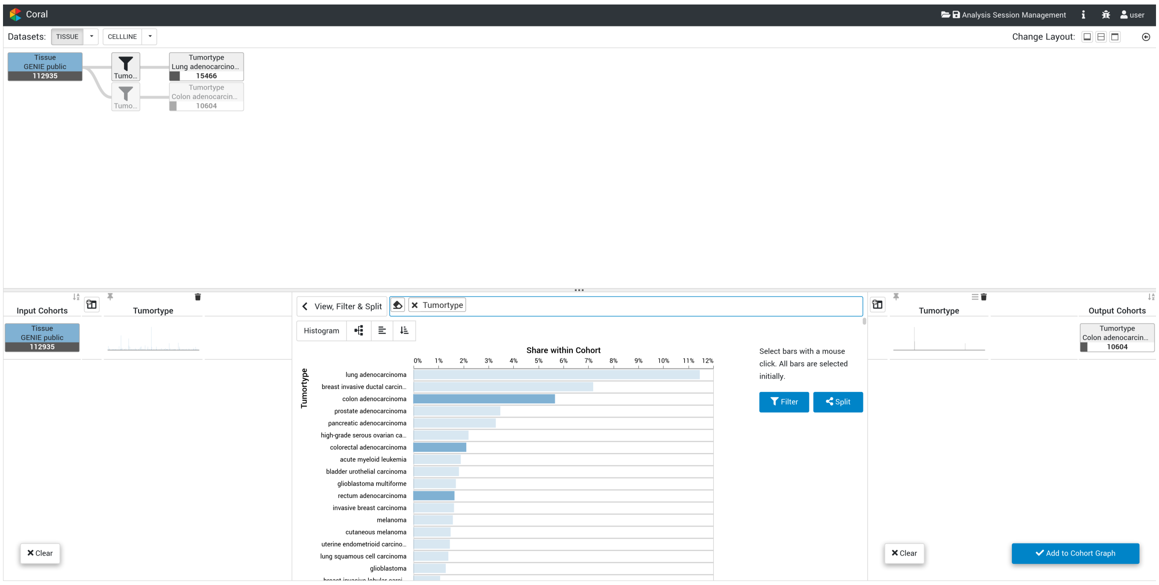

**Supplementary Figure S4**: **Case study 1:** Filtering of the starting cohort (■), which includes all AACR Project GENIE samples, down to colorectal cancer samples. Included tumor types are *colon adenocarcinoma*, *colorectal adenocarcinoma*, and *rectum adenocarcinoma*.

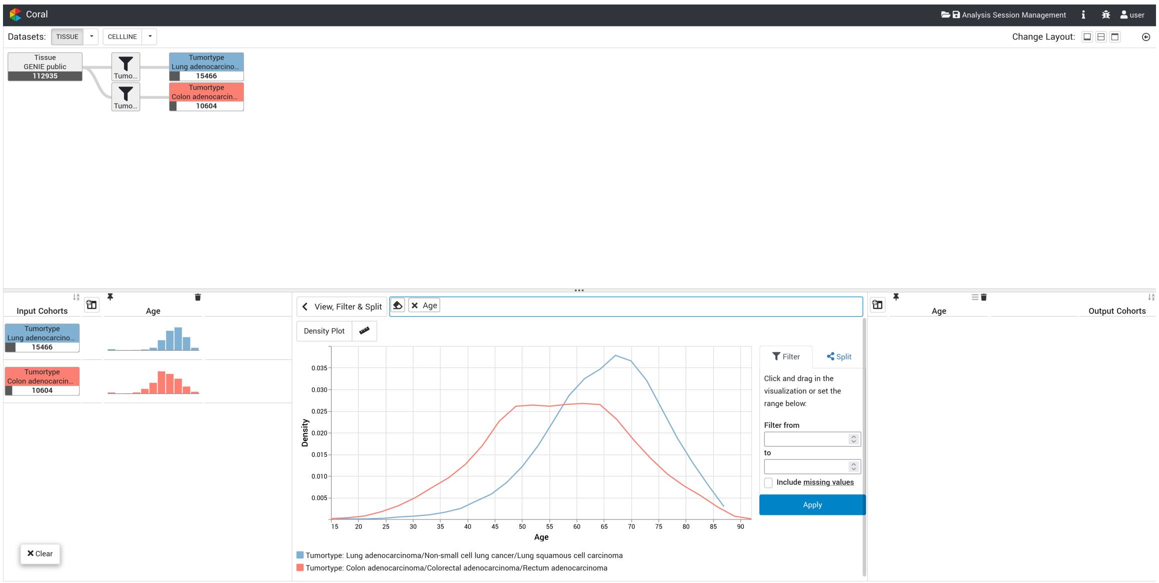

**Supplementary Figure S5**: **Case study 1:** Visualizing the distribution of patient age in the NSCLC (■) and colorectal cancer (■) cohorts.

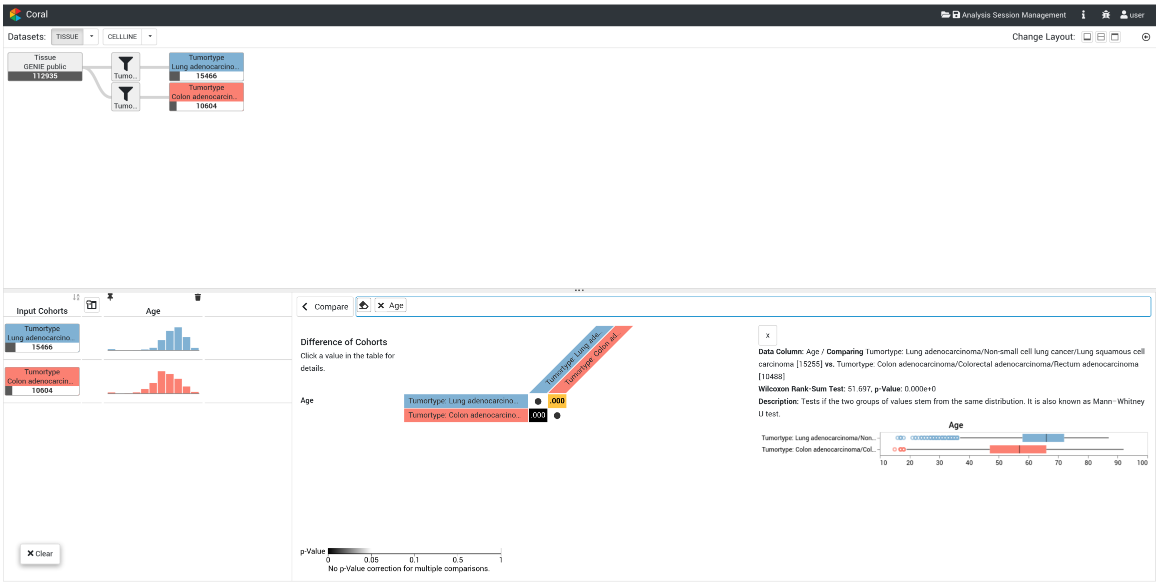

**Supplementary Figure S6**: **Case study 1:** Comparison of the NSCLC (■) and colorectal cancer (■) cohorts in terms of the distribution of patient age. It can be seen that, on average, NSCLC patients are older. This difference is statistically significant.

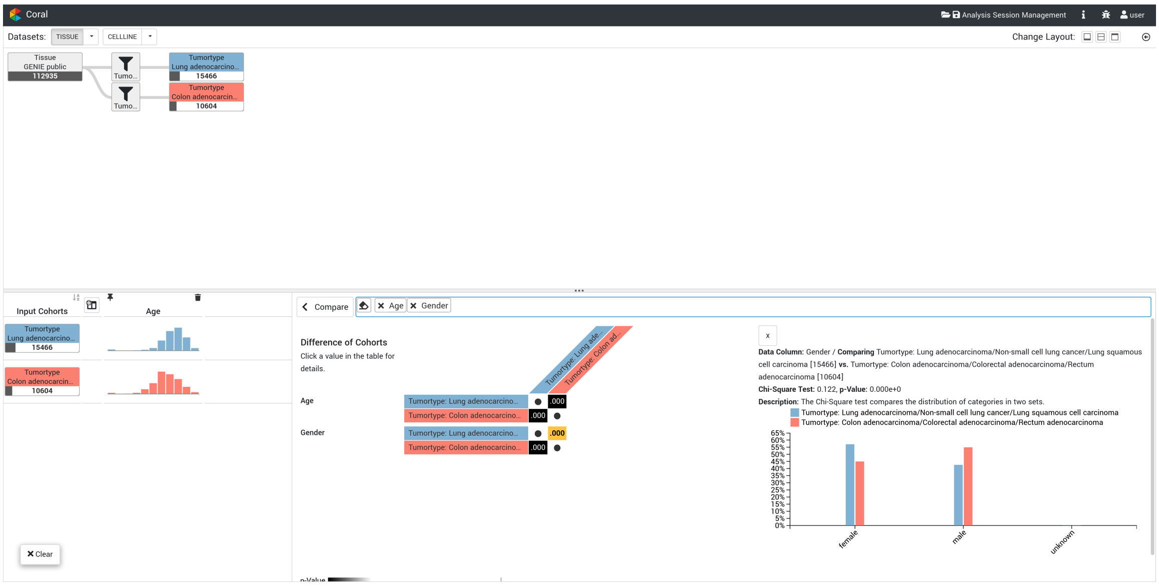

**Supplementary Figure S7**: **Case study 1:** Comparison of the NSCLC (■) and colorectal cancer (■) cohorts in terms of the distribution of patient gender. It can be seen that the NSCLC cohort has a higher proportion of female patients. This difference is statistically significant.

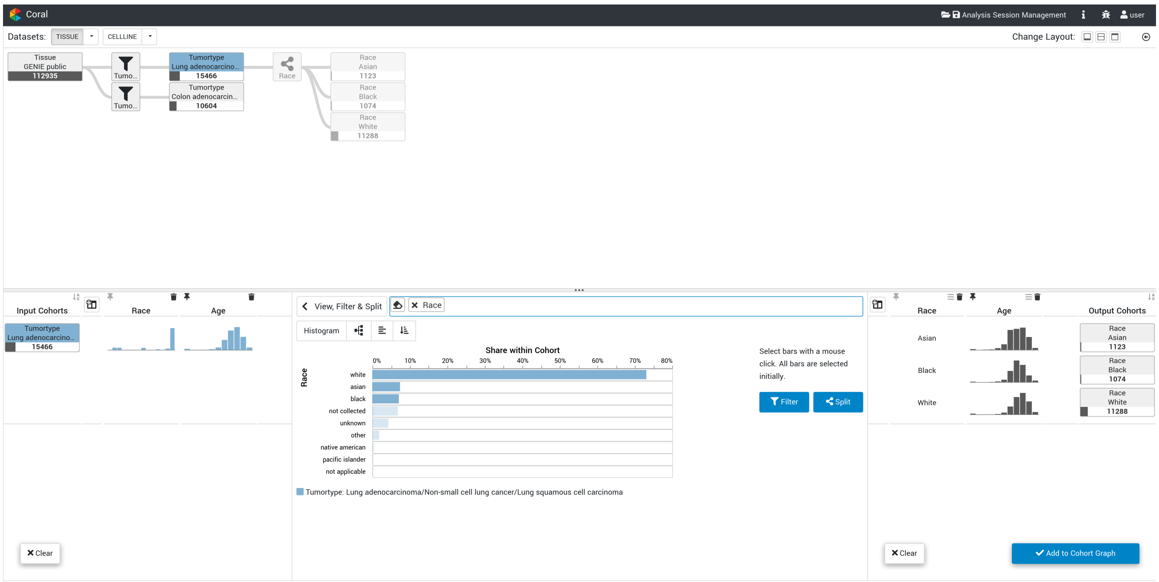

**Supplementary Figure S8**: **Case study 1:** Splitting the NSCLC cohort (■) by the attribute *race* and selecting the three largest populations in the cohort, namely *White*, *Asian*, and *Black*.

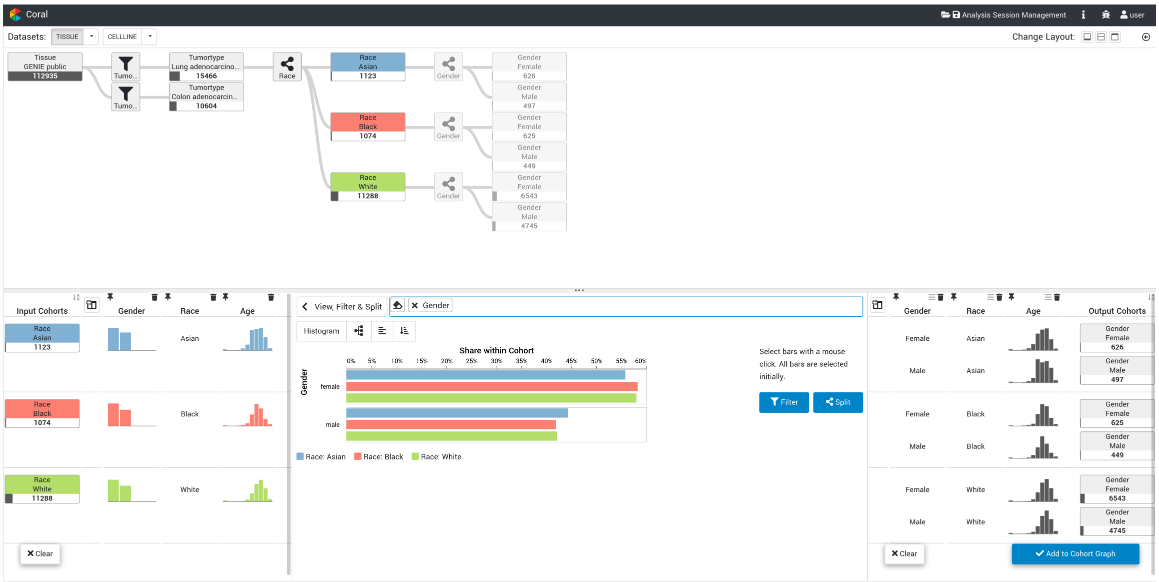

**Supplementary Figure S9**: **Case study 1:** Splitting the three race cohorts (■, ■, ■) by the attribute *gender*.

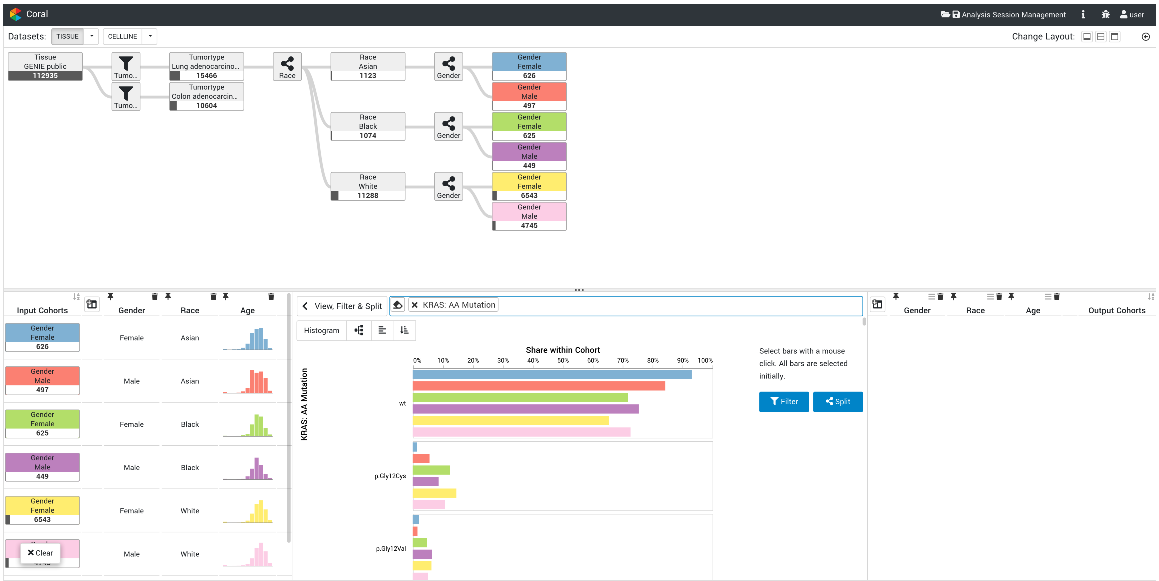

**Supplementary Figure S10**: **Case study 1:** Visualizing the frequencies of various KRAS mutations across the six cohorts generated in the previous steps (■, ■, ■, ■, ■, ■). Link to Coral state in this figure: http://vistories.org/coral-supplementary-figure-10

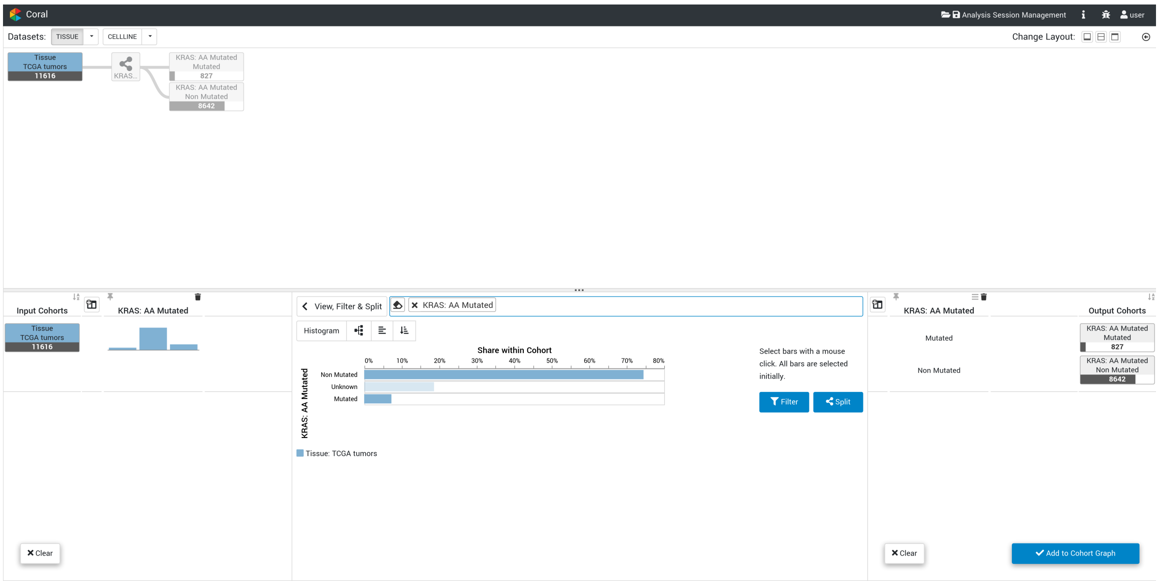

**Supplementary Figure S11**: **Case study 2:** Visualizing the KRAS mutation status distribution for the TCGA tumors dataset (■) and creating two new cohorts—one for *KRAS mutated* and one for *KRAS non-mutated*.

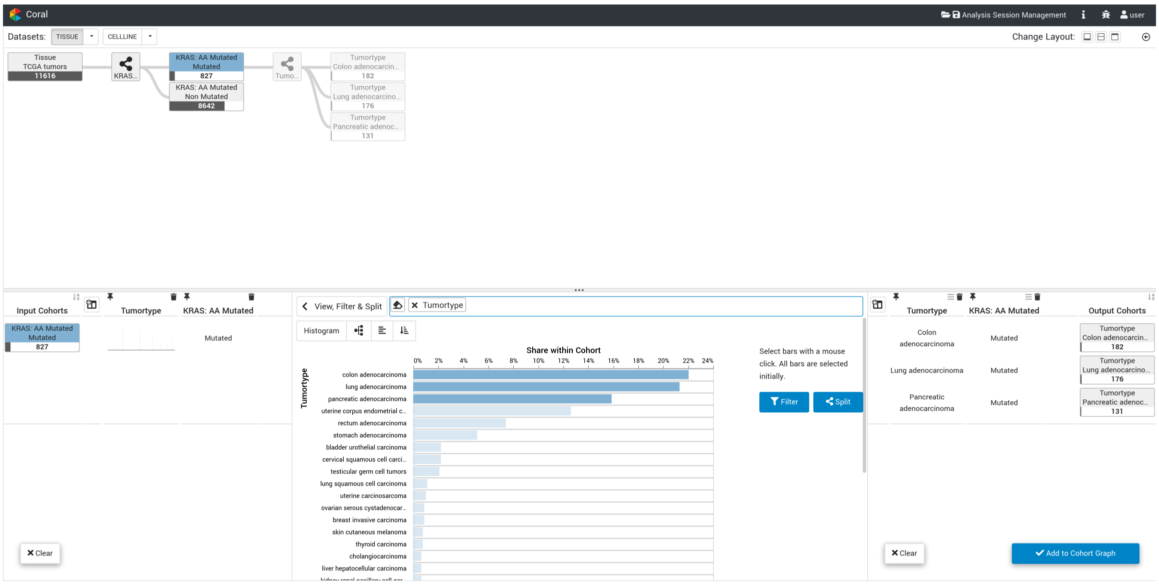

**Supplementary Figure S12**: **Case study 2:** Visualizing the tumor type distribution of the KRAS mutated cohort (■) and creating new cohorts for the three most frequent tumor types—*pancreatic adenocarcinoma*, *lung adenocarcinoma*, and *colon adenocarcinoma*.

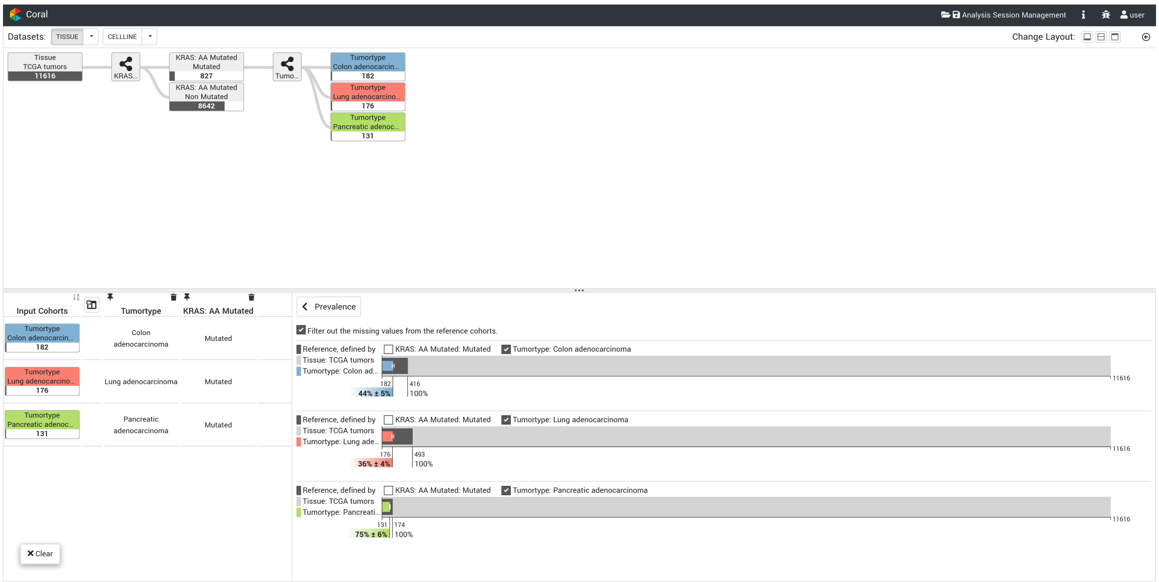

**Supplementary Figure S13**: **Case study 2:** Prevalence of KRAS mutations in three different tumor type cohorts: *colon adenocarcinoma* (■), *lung adenocarcinoma* (■), and *pancreatic adenocarcinoma* (■). It can be seen that KRAS mutations are most prevalent in pancreatic adenocarcinoma patients.

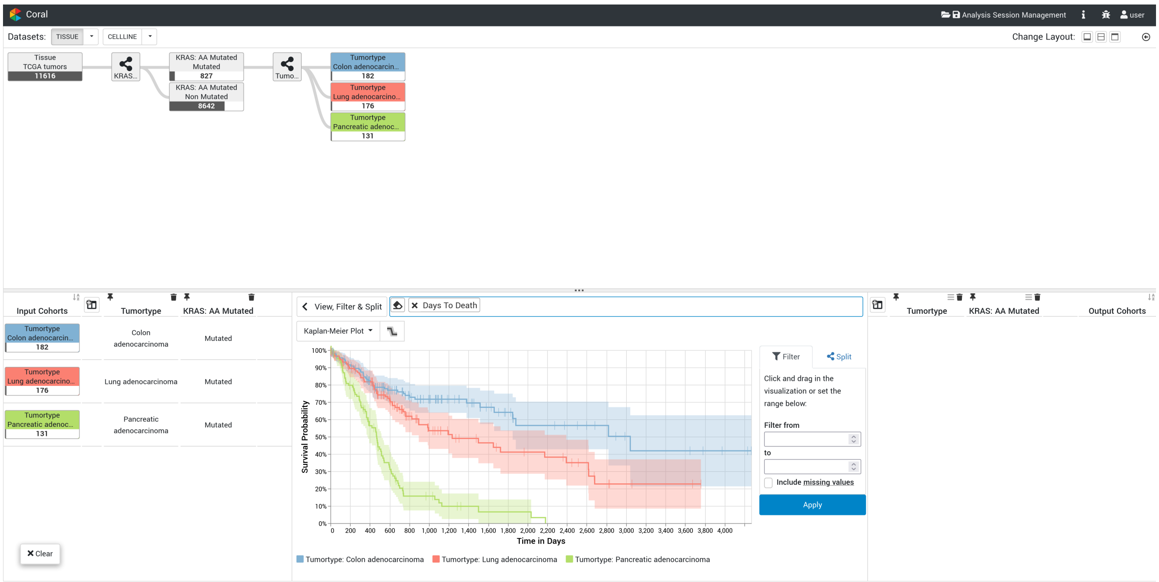

**Supplementary Figure S14**: **Case study 2:** Survival plot of *colon adenocarcinoma* (■), *lung adenocarcinoma* (■), and *pancreatic adenocarcinoma* (■) patients harboring KRAS mutations. It shows that *pancreatic adenocarcinoma* (■) patients have a very poor prognosis.

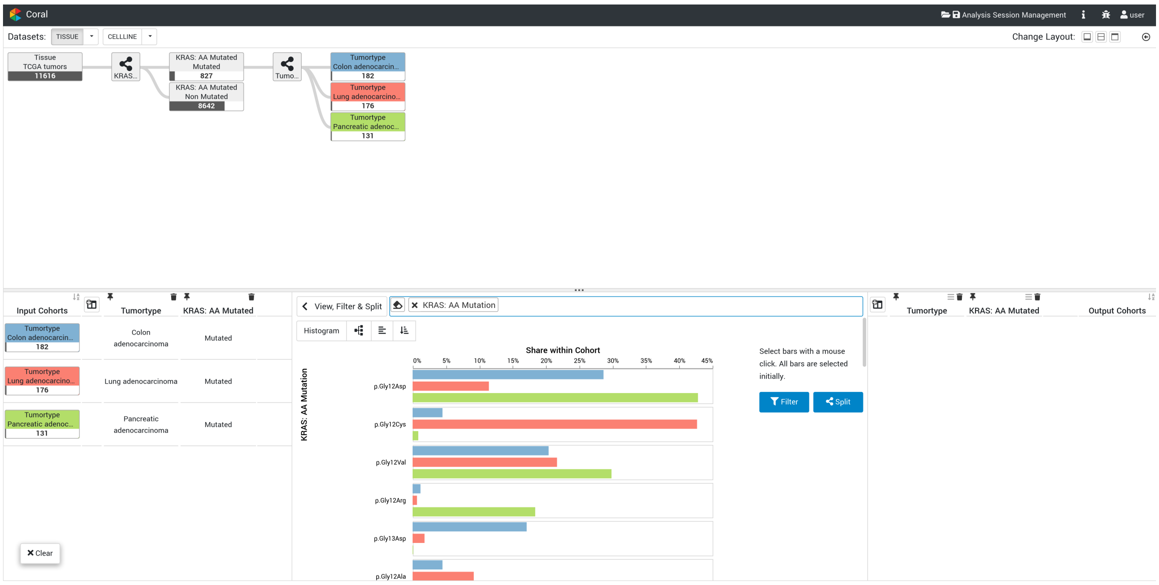

**Supplementary Figure S15**: **Case study 2:** Visualizing the KRAS mutation-type distribution for *colon adenocarcinoma* (■), *lung adenocarcinoma* (■), and *pancreatic adenocarcinoma* (■). The *lung adenocarcinoma* (■) cohort differs significantly from the other two tumor type cohorts in regard to the frequency of specific KRAS mutations.

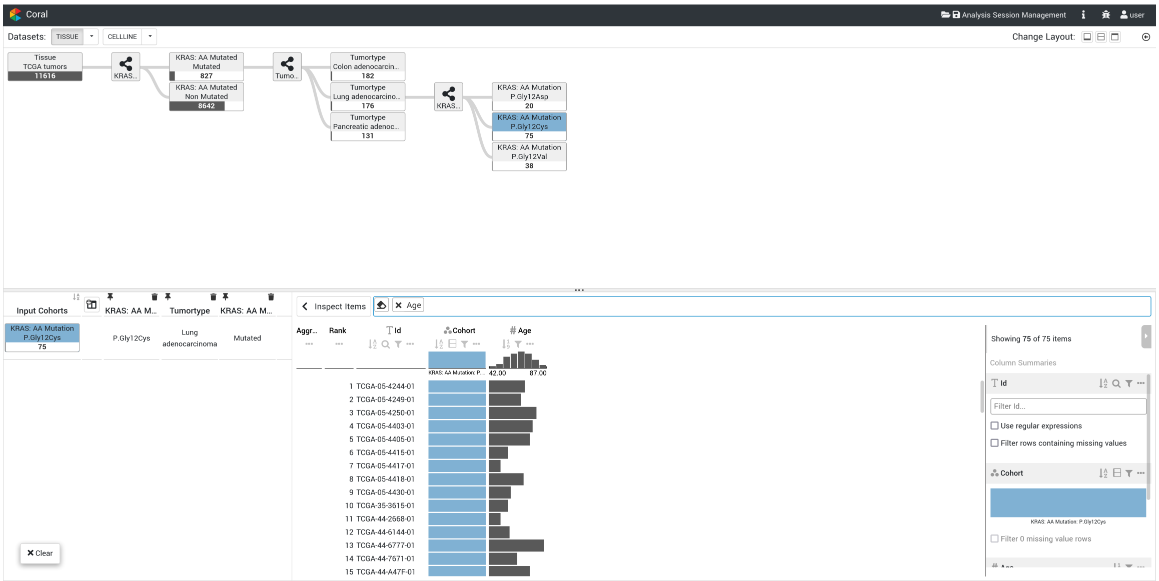

**Supplementary Figure S16**: **Case study 2:** The *Inspect Item* operation shows the individual *lung adenocarcinoma* samples harboring a KRAS Gly12Cys mutation (■).

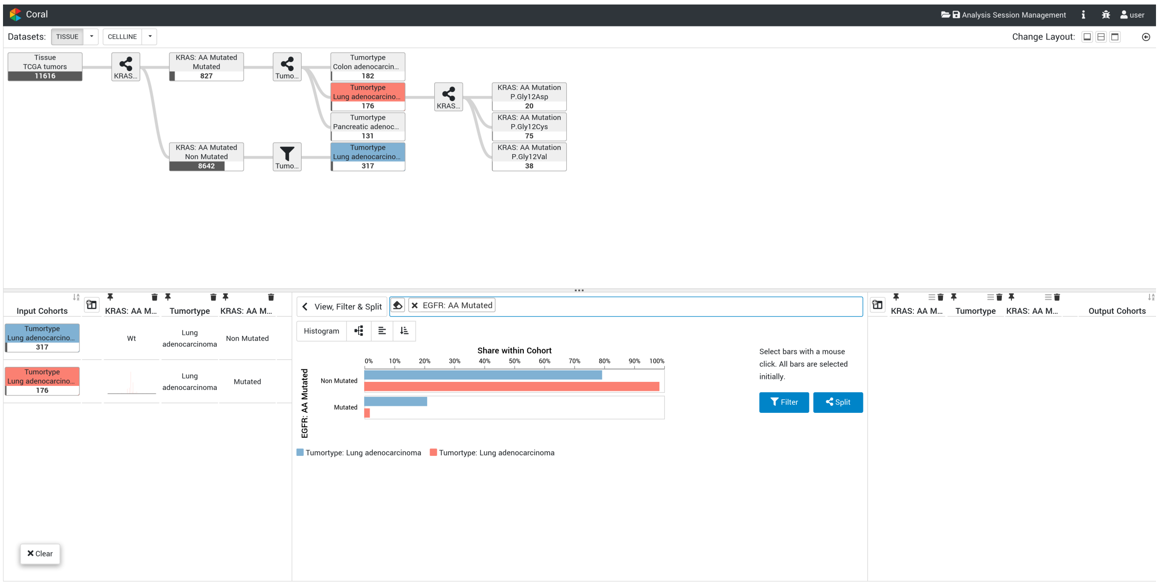

**Supplementary Figure S17**: **Case study 2:** Visualizing the EGFR mutation status for KRAS mutated (■) and KRAS non-mutated (■) *lung adenocarcinoma* samples. It can be seen that EGFR and KRAS mutations are almost mutually exclusive; i.e., they rarely occur together. Link to Coral state in this figure: http://vistories.org/coral-supplementary-figure-17

### 2 Supplementary Tables

| Operation | # of Attributes | Attribute Types | Visualization for One Cohort | Visualization for Multiple Cohorts | Notes |
|---|---|---|---|---|---|
| View, Filter & Split | 1 | Categorical | Bar Chart | Grouped Bar Chart | With absolute or relative scale |
| View, Filter & Split | 1 | Quantitative | Density Plot | Superimposed Density Plot | Area under the curve is not filled, due to higher readability of the superimposed plots |
| View, Filter & Split | 1 | Quantitative | Kaplan-Meier Plot | Superimposed Kaplan-Meier Plot | Kaplan-Meier plot is also known as survival plot, and is used only for attributes related to the survival |
| View, Filter & Split | 2 | Both Categorical | Scatterplot | Superimposed Scatterplot | Dimensionality reduction algorithm used to create the 2D scatterplot |
| View, Filter & Split | 2 | Categorical / Quantitative | Boxplot for each Category | Boxplot for each Category of each Cohort | |
| View, Filter & Split | 2 | Both Quantitative | Scatterplot | Superimposed Scatterplot | |
| View, Filter & Split | 3 or more | Any Combination | Scatterplot | Superimposed Scatterplot | Dimensionality reduction algorithm used to create the 2D scatterplot |
| Prevalence | None | – | Novel Prevalence Encoding | Novel Prevalence Encoding for each Cohort | The novel prevalence encoding allows to flexibly define the reference cohort |
| Inspect Items | Any | Any | Column | Column for each Cohort | Uses Taggle (Furmanova *et al.*, 2020) to show attribute data for each item |
| Compare | Any | Any | Matrix | Matrix | Uses TourDino (Eckelt *et al.*, 2019) to calculate and visualize cohort similarity |

**Supplementary Table S1**: Overview of visualizations used, based on the operation, the number of attributes, attribute types, and their combinations.

### 3 Supplementary Notes

#### 3.1 Terminology

A cohort refers to a group of items, also called records or entities, with shared characteristics. A cohort can be defined based on a single attribute—e.g., patients of a certain age group—or on multiple attributes—e.g., elderly lung cancer patients with a *KRAS* mutation.

The basis for forming cohorts are datasets in the form of multi-attribute tabular data. We refer to the columns of a table as attributes, and the rows as items (Munzner, 2014, p. 25). Within a column, all values are of the same type: quantitative or categorical.

Our public instance of Coral is pre-loaded with multiple genomics datasets, described in Section 3.7.

#### 3.2 User Goals

A cohort analysis tool needs to serve two high-level purposes. Users can either aim to generate new hypotheses, such as the identification of factors that contribute to the development, progression, or spread of a disease, or to verify existing hypotheses, like confirming the effect of a specific mutation on a tumor type. In this section, we introduce the specific operations users need to be able to fulfill as part of a cohort analysis.

We decided to structure the operations by means of three operation categories: (1) cohort creation, (2) cohort characterization, and (3) cohort tracking. Operations from these categories are not carried out in a sequence but users want to be able to switch between them as needed during the analysis. The cohort creation operations are *Filter & Split*. The *View* operation, *Compare* operation, *Prevalence* operation, and *Inspect Items* operation are part of the cohort characterization. And the *Cohort Evolution View* keeps track of all cohorts.

#### 3.3 Related Tools

##### 3.3.1 cBioPortal

cBioPortal (Cerami *et al.*, 2012; Gao *et al.*, 2013) is an interactive web application that allows researchers to access and analyze complex genomic data of different cancer genomics projects. It is a well established application in the field of cancer research.

cBioPortal, like Coral, allows the definition and comparison of sample groups (cohorts), however, both applications were developed with different use cases in mind. The main focus of cBioPortal lies on the analysis of selected genes, the analysis of individual patients, and the exploration of studies. In contrast, Coral focuses more on the iterative cohort creation, their analysis and comparison. Due to this difference, cBioPortal has some shortcomings in regard to cohort creation, as outlined below.

There are several limitations regarding the definition of cohorts based on mutation data. For instance, creating cohorts with specific mutations (e.g., *KRAS G12C*) has to be done via OQL (Onco Query Language; http://www.cbioportal.org/oql), which is a cBioPortal specific language that makes it difficult for users new to cBioPortal to use all its functionalities. Additionally, defining cohorts based on the absence of one or multiple mutations (e.g., samples that are *EGFR* wild-type) is not trivial. As shown in the cBioPortal group comparison tutorial (https://www.cbioportal.org/tutorials#group-comparison) this can only be achieved by defining a 'baseline cohort' and then using non-overlapping sets to define the actual cohorts. This cohort creation process, utilizing auxiliary cohorts, can be too complicated for non-experienced users. More importantly, this approach assumes that every gene that is not mutated is wildtype, but this assumption does not hold true for data sources like the AACR Project GENIE which constitutes a heterogeneous mix of gene panels, i.e. not the exact same set of genes is probed on each panel.

Another limitation is that the iterative process to define a larger set of cohorts can be cumbersome. For example, creating cohorts by multiple subsequent split operations (e.g., split by ethnicity followed by split by gender) is not directly supported. Furthermore, tracking the provenance of all generated cohorts is not possible in cBioPortal. This makes it difficult to keep the overview of the cohorts and limits the reproducibility of the performed analysis.

##### 3.3.2 Other Tools

There are many tools, besides cBioPortal, that allow the creation and analysis of cohorts. Most of them are designed for specific datasets and analysis goals. For example, Composer by Rogers *et al.* (2019) focuses on the treatment of patients with lower back conditions. The application allows the analyst to create cohorts and then to decide which treatment would result in the best outcome. Other cohort analysis tools are, for instance, CAVA by Zhang *et al.* (2015) and a prostate cancer analysis tool by Bernard *et al.* (2015). All of these three applications allow users to create cohorts, however, refinement of multiple cohorts at once is not possible. Furthermore, they all have some kind of history functionality to show which filters were applied, but none of them show how the created cohorts relate to each other. Additionally, it is not possible to visualize data of multiple cohorts in these tools. The only exception is Composer that visualizes the main attribute of interest of multiple cohorts. This makes it cumbersome to compare the different cohorts with each other.

#### 3.4 Session Management: Reproducing and Sharing

An important consideration in the field of (bio)medicine is being able to reproduce an analysis. This reproducibility is achieved through a session manager that allows users to save, share, and revisit their analysis session. As a consequence, users need to log in before they can use Coral; however, to get around the registration process for new users, the publicly available Coral instance auto-generates accounts.

During an analysis session, the operations performed by the user are recorded and stored in the local cache of the browser. A user can then decide to make an analysis session persistent, which moves it from the browser storage to a database on the Coral server. A persistent session can be made public and shared by copying the URL shown in the browser. When a persistent session is loaded, Coral restores the cohorts of the analysis, including the operations that were used (Gratzl *et al.*, 2016). In **Supplementary Figure S10** and **Supplementary Figure S17** we include a link in the figure caption with which the cohorts can be reproduced.

#### 3.5 Workflow

The Coral workflow consists of four main steps, as illustrated in **Supplementary Figure S1**: (1) select cohorts, (2) select operations that will be applied to the cohorts, (3) define the output cohorts, and (4) adding these output cohorts to the *Cohort Evolution View.*

##### 3.5.1 Step 1: Select Cohorts

The user starts the analysis by selecting one or more cohorts from the *Cohort Evolution View.* The selection of a cohort adds it to the *Input Area* of the *Action View* and assigns a color to the cohort that is used consistently in all visualizations. We refer to these cohorts as the *input cohorts* as they serve as the inputs for the creation and characterization operations (see **Supplementary Figure S1❶**).

##### 3.5.2 Step 2: Select Operation

In the *Operation Area*, two types of operations are available: creation and characterization. Characterization operations give insights into the input cohorts but do not manipulate them. They, therefore, have no output. Users can choose a different operation or use the *Cohort Evolution View* to switch inputs when they are done characterizing the input cohorts (see **Supplementary Figure S1❷** and **Supplementary Figure S2**). Creation operations allow users to create new cohorts based on different attributes and attribute combinations. For some operations, e.g. Filter & Split, users have to select one or more attributes of interest with the *Search Bar* at the top of the *Operation Area.* This search bar, as the name indicates, can be used to browse through a list of available attributes or search for specific ones. **Supplementary Table S1** lists the offered visualizations for the different operations, attribute types, and attribute combinations.

###### 3.5.3 Step 3: Define Output Cohorts

Creation operations—Filter & Split—output new cohorts, shown in the *Output Area.* The Split operation allows splitting an input cohort into multiple output cohorts. Whereas the Filter operation creates an output cohort from a subset of the input cohort. (see **Supplementary Figure S1❸**). Users can select categories or value ranges to filter or split the input cohorts. The *Input Area* and the *Output Area* both use a tabular layout that shows input cohorts, their related output cohorts, and attribute distributions.

###### 3.5.4 Step 4: Add Output Cohorts to Evolution View

The output cohorts and the operations applied are displayed as a preview in the *Cohort Evolution View.* Before confirming the set of cohorts to be finally added, users can deselect empty cohorts or cohorts of no interest. This additional confirmation step avoids cluttering the *Cohort Evolution View* during data exploration (see **Supplementary Figure S1❹**).

##### 3.6 Implementation

Coral is publicly available at `https://coral.caleydoapp.org/` and its source code is available at `https://github.com/Caleydo/coral`. The web-client is implemented in TypeScript and utilizes Vega-Lite (Satyanarayan *et al.*, 2017) to create visualizations. The server-side uses Python to provide the REST interface between front-end and back-end as well as the communication with a Postgres database. The public version is deployed on Amazon Web Services (AWS).

##### 3.7 Data Processing & Integration

The publicly deployed Coral instance contains the following datasets:

- AACR Project GENIE public release version 9.0 (AACR Project GENIE Consortium, 2017) `https://www.aacr.org/professionals/research/aacr-project-genie/`
- The Cancer Genome Atlas (TCGA) `https://cancergenome.nih.gov`
- Cancer Cell Line Encyclopedia (CCLE, Barretina *et al.* (2012)) `https://portals.broadinstitute.org/ccle`
- Project DRIVE (McDonald *et al.*, 2017)
- Avana CERES (Meyers *et al.*, 2017)

###### 3.7.1 AACR Project GENIE

The American Association of Cancer Research (AACR) Project GENIE is a public cancer registry of real-world data assembled through data sharing between 19 of the leading cancer centers in the world. GENIE stands for *Genomics Evidence Neoplasia Information Exchange* and its goal is to power precision oncology and clinical decision making (`https://www.aacr.org/professionals/research/aacr-project-genie`).

We downloaded the latest public release v9.0-public on February 8th, 2021 from `http://synapse.org/genie`. We converted the mutation file (data_mutations_extended.txt) into separate VCF files per sample and annotated them using Ensembl VEP v70. Upon insertion into the database, which is based on hg38, we stripped the genomic coordinates of the mutations and only inserted the translated cDNA and amino acid changes. We checked that the transcript structures of genes listed on any of the gene panels were comparable between hg19 and hg38.

###### 3.7.2 TCGA Sample Selection

The TCGA sample cohorts COADREAD, FPPP, GBMLGG, KIPAN, and STES were excluded since these are combined or experimental cohorts.

##### 3.7.3 TCGA Metadata

The R package TCGAbiolinks (Version 2.5.9, Colaprico *et al.* (2016)) was used to extract sample and patient information for TCGA samples by using a custom-made R script.

##### 3.7.4 TCGA Gene Expression Data

The GDC Data Portal's interface (`https://portal.gdc.cancer.gov/`) was used to generate a manifest file of all data files that mapped the fields "Program" = "TCGA", "Data Type" = "Aligned Reads", "Experimental Strategy" = "RNA-Seq", and "Workflow Type" = "STAR 2-Pass". Using the GDC Data Transfer Tool, the data was transferred and pre-processed via the commands samtools (Li *et al.*, 2009) collate and samtools fastq to ultimately generate FASTQ files, containing the unmapped reads. All samples were subsequently processed with a harmonized RNA-seq pipeline, described in Hofmann *et al.* (2021).

##### 3.7.5 TCGA Mutation Data

We downloaded the Mutect2 VCF files from the Genomics Data Commons (`https://docs.gdc.cancer.gov/Data/Bioinformatics_Pipelines/DNA_Seq_Variant_Calling_Pipeline/`) in February 2018. We reannotated the VCF files using a combination of Ensembl VEP v86 and v93 focusing on mutations that pass all filtering criteria as employed by the TCGA Mutect2 pipeline (labeled as PASS). Mutations that failed the filtering but were recurrently present in the COSMIC database were added.

##### 3.7.6 TCGA Copy Number Data

TCGA SNP6 copy number segmentation data was downloaded from NIH GDC (`https://portal.gdc.cancer.gov/`, Grossman *et al.* (2016)) on December 3, 2018. The segmentation information was obtained from the files `*nocnv_grch38.seg.v2.txt`. Gene-wise copy numbers were determined by overlapping the segmentation information with Ensembl v86 gene annotation. If a gene was covered by a single segment, the copy number of the segment was assigned to the gene. If a gene was covered by multiple segments, a weighted average copy number was computed based on the size of the overlap between the gene and each segment. Relative copy numbers <= 1.0 were considered as "deep deletion", and relative copy numbers >= 3.5 were considered as "amplification".

##### 3.7.7 CCLE Metadata

Cell line names and descriptions (organ of origin, metastatic site, histology type, morphology, growth type, gender, and age at surgery) were taken from the provider's cell-line data sheet. If a cell line was available from various vendors, the cell-line name was taken from the top rank in a hierarchy of vendors in the following order: atcc, dsmz, ecacc, jcrb, iclc, riken, kclb.

##### 3.7.8 CCLE Gene Expression Data

Raw FASTQ data for all CCLE cell lines published in Ghandi *et al.* (2019), were downloaded via the European Nucleotide Archive (accession number PRJNA523380). All data were processed identically to the TCGA data as described above.

##### 3.7.9 CCLE Mutation Data

Variant calling of cell lines followed community best practices. The reads were aligned using BWA version 0.7.17 against the reference genome hg38. We used Picard 2.17.8 to remove duplicates and strelka2 2.8.4 to call somatic mutations. We used an unmatched normal sample from the 1000G project (NA12878) as the normal sample. Called mutations were annotated by a combination of Ensembl VEP v86 and v93, flagging putative germline variation by using population frequencies from the 1000G project and gnomAD. Putative alignment artifacts were filtered out using a mutation blacklist derived from the Sanger COSMIC Cell line Project VCF files, for which putative artifacts/germline variation is flagged in the VCF files. We computed coverage statistics for each gene in each sample: In the absence of a mutation, we called a gene wild-type if and only if at least 80% of bases of the gene body (excluding the first exon) were sufficiently covered, and NA otherwise.

##### 3.7.10 CCLE Copy Number Data

SNP6 CEL files were downloaded from `https://cghub.ucsc.edu/` in October 2012. Relative copy number segments were computed using the R package `aroma.affymetrix` version 3.1.0 (Bengtsson *et al.*, 2008a,b, 2009) and Rawcopy version 1.1 (Mayrhofer *et al.*, 2016): the SNP6 data was processed with the AROMA method CRMA v2, where the 50 samples with the least amount of copy number alterations based on Rawcopy were used to calculate the reference intensities. This was followed by CBS segmentation. Afterward, the copy number segments were overlapped with Ensembl v86 gene annotation analogously to the TCGA processing in order to obtain gene-wise relative copy number values. "Amplification" and "deep deletion" status were also assigned as in the TCGA processing. Absolute copy number segments were computed using PICNIC version c_release 2010-10-29 (Greenman *et al.*, 2010) with reference files adapted for reference genome hg38 and default parameters. The resulting segments were overlapped with Ensembl v86 gene annotation as in the TCGA processing in order to obtain gene-wise absolute copy number values.

##### 3.7.11 DRIVE Data

DRIVE (deep RNAi interrogation of viability effects in cancer) is a large shRNA screen of ∼8000 genes and ∼400 cancer cell lines (McDonald *et al.*, 2017). Raw data and processed RSA and ATARIS scores were transferred via email by the authors. siRNAs targeting multiple genes were discarded. Gene symbols were translated into Ensembl stable identifiers for genes by using the official gene symbol provided by the Ensembl database Version 86. Cell-line names are identical to CCLE cell-line names and were translated to the Boehringer Ingelheim cell-line nomenclature.

##### 3.7.12 Avana Data

The Avana single guide RNA (sgRNA) library was used in a large CRISPR/Cas9 loss-of-function screen (Meyers *et al.*, 2017) of ∼770 cell lines and ∼17,500 genes (version 21Q1). Processed unscaled CERES scores (representing the estimated gene knockout effects) were taken from `https://depmap.org` and the final CERES scores were calculated (using the common essentials and non-essential genes from the same source) based on the original method published in Meyers *et al.* (2017). Entrez Gene IDs were translated into Ensembl gene identifiers using the Ensembl gene database Version 86. As for the DRIVE dataset, CCLE cell-line names were used to translate cell-line identifiers into Boehringer Ingelheim's cell-line names.

#### 3.8 Case Study 1

The main focus of this case study are $KRAS^{G12C}$ somatic mutations. The gene *KRAS* is mutated in ∼15% of all human cancers making it one of the most frequently mutated cancer-causing genes. *G12C* (a glycine-to-cysteine substitution at codon 12) is one of the most prevalent *KRAS* mutations and the target of several new drugs in clinical development.

Here, we reproduce some of the findings from a recently published article about $KRAS^{G12C}$ mutations (Nassar *et al.*, 2021). The authors leverage the AACR Project GENIE patient cohort to assess race and gender differences with respect to $KRAS^{G12C}$ mutation frequencies in Non-Small Cell Lung Cancer (NSCLC) and colorectal cancer patients.

We start the analysis by loading the AACR Project GENIE public dataset. We then create two sub-cohorts containing the NSCLC and colorectal cancer samples. To do so, we first filter the GENIE cohort using the tumor types *lung adenocarcinoma*, *lung squamous cell carcinoma*, and *non-small cell lung cancer* which comprise the major subtypes of NSCLC (**Supplementary Figure S3**). Next, we filter the GENIE cohort using the tumor types *colon adenocarcinoma*, *colorectal adenocarcinoma*, and *rectum adenocarcinoma* which together constitute colorectal cancers (**Supplementary Figure S4**).

To get an initial impression of the demographics we select both cohorts and use the *View* operation to visualize the distribution of the numerical attribute *age* (**Supplementary Figure S5**). Additionally, we add the *age* attribute to the cohort tables in the *Input* and *Output Area*. We see that NSCLC patients in the GENIE dataset seem to be, on average, older compared to the colorectal cancer patients. To confirm that this difference is statistically significant, we use the *Compare* operation (**Supplementary Figure S6**). Additionally, we investigate potential differences regarding the gender distribution and see that the NSCLC cohort has a significantly higher proportion of female patients (**Supplementary Figure S7**).

Subsequently, we want to check potential race and gender differences in the data with respect to the $KRAS^{G12C}$ mutation frequency. For this analysis we focus on NSCLC patients; we can apply the analysis steps, however, in the same way to the colorectal cancer cohort. We perform the *Split* operation to the NSCLC cohort using the categorical attribute *race*. We then create cohorts for the three largest populations in the dataset, namely *White*, *Asian*, and *Black* (**Supplementary Figure S8**). Next, we generate six new cohorts via the *split* operation on the categorical attribute *gender* (**Supplementary Figure S9**). Finally, we use the *View* operation to visualize the frequency of the different *KRAS* mutations in each of the cohorts using the categorical attribute *KRAS: Amino Acid Mutation* (**Supplementary Figure S10**).

$KRAS^{G12C}$ mutations (here called *Gly12Cys*) in NSCLC are less frequent in Asian women (∼1%) compared to Asian men (∼5%), but more frequent in Black and White women (∼12% and ∼14%) compared to Black and White men (∼9% and ∼11%), respectively. Furthermore, the prevalence of this mutation is overall lower for Asian compared to Black and White. These observations match the findings of Nassar *et al.* (2021) very well (compare the **Supplementary Figure S10** *Gly12Cys* bar chart with Fig. 1B of Nassar *et al.* (2021)). Minor differences in the numbers are mainly caused by the use of different GENIE releases (v8.0 in Nassar *et al.* (2021) versus v9.0 in Coral).

The visualization in Coral easily allows a closer investigation of other *KRAS* mutations. We can see, for instance, that there is a lower gender difference for *G12V* (*Gly12Val*) mutations in Asians. Furthermore, the user can easily select the samples that harbor the mutation of interest, filter for them, and then analyze them in even further detail.

##### 3.9 Case Study 2

This case study summarizes an analysis session carried out by a collaborator with a background in bioinformatics. We demonstrate how Coral helps and supports the analysis process visually to get to the results of the session.

Coral provides a cancer genomics dataset with tissue sample data collected by the The Cancer Genome Atlas (TCGA; `https://cancergenome.nih.gov`) project for 33 cancer types. The dataset contains meta-data, such as the age and gender of the patients, and gene expression, mutation, and copy number data of the samples.

The main focus of this analysis session is the gene *KRAS*. As indicated above, it is mutated in ∼15% of all human cancers making it one of the most frequently mutated cancer-causing genes. Despite several decades of research into this gene, there are still no approved drugs available that target *KRAS*. However, several new drugs are currently in clinical development. The aim of this analysis session is to investigate the landscape of *KRAS* mutations across different types of cancer and to understand which patient populations could benefit from the new drugs.

The analyst starts by selecting the TCGA tumors dataset and then uses the *Split* operation to split the cohort into *KRAS* mutated and *KRAS* non-mutated cohorts (**Supplementary Figure S11**).

Afterward, he assesses the distribution of tumor types in the *KRAS* mutated cohort and observes that *pancreatic adenocarcinoma*, *lung adenocarcinoma*, and *colon adenocarcinoma* are the most frequent tumor types of the TCGA *KRAS* mutated cases. The analyst selects these and creates the corresponding cohorts using the *Split* operation (**Supplementary Figure S12**).

Subsequently, he assesses the *KRAS* mutation prevalence in these three tumor types and can confirm the expectation that *KRAS* is very frequently mutated in these patients (∼75% in pancreatic, ∼44% in colon, and ∼36% in lung adenocarcinoma), as shown in **Supplementary Figure S13**. Furthermore, assessing the survival data of these cohorts (**Supplementary Figure S14**) shows that in particular pancreatic adenocarcinoma patients typically have a very poor prognosis. This highlights the importance of developing drugs against mutated *KRAS*.

As the next step, the analyst investigates which specific *KRAS* mutations the patients have. He selects the three cohorts and visualizes the attribute *KRAS: Amino Acid Mutation*. From the data it becomes very obvious that lung adenocarcinomas differ significantly from the other two tumor types with respect to the frequency of specific *KRAS* mutations. The three most common mutations in lung adenocarcinomas are *Gly12Cys*, *Gly12Val*, and *Gly12Asp* (**Supplementary Figure S15**). In contrast, the three most common mutations in pancreatic and colon adenocarcinomas are *Gly12Asp*,

*Gly12Val*, and *Gly12Arg*, and *Gly12Asp*, *Gly12Val*, and *Gly13Asp*, respectively. The *KRAS* drugs that are most advanced in clinical development target the mutation *Gly12Cys*. So, this analysis shows that lung adenocarcinoma patients would benefit most from them due to the large fraction of patients with that specific *KRAS* mutation.

To learn more about these lung adenocarcinoma patients, the analyst selects the top three *KRAS* mutations and splits the cohort by them. Selecting the *Gly12Cys* cohort and opening the *Inspect Items* operation reveals the individual samples that could have benefited from the treatment mentioned above. This list also allows adding further information for each patient, for example, its age (**Supplementary Figure S16**).

As the last aspect of the investigation, the analyst is interested in the mutation frequency of the gene *EGFR* in lung adenocarcinoma patients with and without *KRAS* mutations. *EGFR* is another important cancer gene, especially in the context of lung cancer; and several drugs targeting it have been approved. The analyst selects the cohort of *KRAS* non-mutated samples and filters them by the tumor type lung adenocarcinoma. Afterward, he additionally selects the lung adenocarcinoma cohort for the *KRAS* mutated samples generated in one of the previous steps and visualizes the frequency of *EGFR* mutations. The analyst observes that ∼20% of lung adenocarcinoma patients without *KRAS* mutation harbor an *EGFR* mutation (and could therefore potentially benefit from related treatments), whereas almost none of the selected *KRAS* mutated samples have a mutation in *EGFR* (**Supplementary Figure S17**).

So, mutations in *KRAS* and mutations in *EGFR* seem to be mutually exclusive. This can be explained by the fact that both genes belong to the same signaling pathway and that over-activation of one is sufficient for tumor development.

The results of this analysis session show that *KRAS* is mutated in many cancers, especially pancreatic, colon, and lung adenocarcinomas, making them the main target populations for *KRAS* drugs in general. But it also highlights that there is a clear difference between these tumor types with respect to the kind of *KRAS* mutations that are present. This has a big influence on the available treatment options. For instance, drugs targeting *KRAS Gly12Cys* are most relevant for lung adenocarcinoma. Furthermore, the analysis shows that mutations in *KRAS* and *EGFR* usually occur in a mutually exclusive manner.

In this session, the analyst shows that switching between creating and characterizing cohorts is done without interfering with the analysis process, which simplifies and speeds-up the analysis. Furthermore, being able to handle many different cohorts simultaneously allows one to easily manage even more complex use cases. Finally, the availability of the Prevalence operation is particularly useful for disease related analyses like this one, since it allows to perform prevalence estimations without the need of defining reference cohorts manually.

#### 3.10 Future Work

To further extend the utility of Coral, we identified the following aspects for future work.

**Session management.** The current session management of Coral stores the creation of cohorts. We plan on extending the stored sessions with further analysis steps, i.e., which attribute was visualized. This would allow the user to better understand why and how certain cohorts were created in the analysis session.

**Extending data and metadata.** To further improve the utility of Coral for cancer research, we plan to extend the included data and metadata. This includes adding additional data sets as well as providing more detailed information and assessments related to the already available data, as, for instance, classifying mutations into *drivers* and *passengers / variants of unknown significance (VUS)*.

**Support for longitudinal data.** Coral currently does not support longitudinal data (e.g., data from multiple samples of one patient at different time points or a detailed treatment history). Currently, these kinds of data sets are still rare. However, we expect this type of data to become more prevalent in the future and therefore plan to extend Coral to support such use cases.

**Integration with existing applications.** To further support a wider range of tasks without re-implementing functionality from other established tools, we plan to investigate the possibility of making Coral interact with existing tools. In particular, we plan to explore ways of tightly integrating Coral with cBioPortal.

**Going beyond cancer research.** Finally, the current focus of Coral and its database is on cancer genomics. However, technically and conceptually, Coral can be applied to data and problems from other fields, for instance, to analyze the data of COVID-19 patients of a country or students of a university. Therefore, we plan to support the upload of custom data and connections to additional databases in the future. By doing so, Coral will be able to cater the needs of an even broader range of researchers in other disciplines.

### Supplementary References

AACR Project GENIE Consortium (2017). AACR Project GENIE: powering precision medicine through an international consortium. *Cancer discovery*, **7**(8), 818–831.

Barretina, J. *et al.* (2012). The Cancer Cell Line Encyclopedia enables predictive modelling of anticancer drug sensitivity. *Nature*, **483**(7391), 603–607.

Bengtsson, H. *et al.* (2008a). aroma.affymetrix: A generic framework in R for analyzing small to very large Affymetrix data sets in bounded memory. Technical Report 745, Department of Statistics, University of California, Berkeley.

Bengtsson, H. *et al.* (2008b). Estimation and assessment of raw copy numbers at the single locus level. *Bioinformatics*, **24**, 759–767.

Bengtsson, H. *et al.* (2009). A single-array preprocessing method for estimating full-resolution raw copy numbers from all Affymetrix genotyping arrays including GenomeWideSNP 5 & 6. *Bioinformatics*, **25**, 2149–2156.

Bernard, J. *et al.* (2015). A Visual-Interactive System for Prostate Cancer Cohort Analysis. *IEEE Computer Graphics and Applications*, **35**(3), 44–55.

Cerami, E. *et al.* (2012). The cBio Cancer Genomics Portal: An Open Platform for Exploring Multidimensional Cancer Genomics Data. *Cancer Discovery*, **2**(5), 401–404.

Colaprico, A. *et al.* (2016). TCGAbiolinks: an R/Bioconductor package for integrative analysis of TCGA data. *Nucleic acids research*, **44**(8), e71.

Eckelt, K. *et al.* (2019). TourDino: A Support View for Confirming Patterns in Tabular Data. In *Proceedings of the EuroVis Workshop on Visual Analytics*, pages 7–11.

Furmanova, K. *et al.* (2020). Taggle: Combining overview and details in tabular data visualizations. *Information Visualization*, **19**(2), 114–136.

Gao, J. *et al.* (2013). Integrative Analysis of Complex Cancer Genomics and Clinical Profiles Using the cBioPortal. *Science Signaling*, **6**(269), pl1.

Ghandi, M. *et al.* (2019). Next-generation characterization of the Cancer Cell Line Encyclopedia. *Nature*, **569**(7757), 503–508.

Gratzl, S. *et al.* (2016). From Visual Exploration to Storytelling and Back Again. *Computer Graphics Forum*, **35**(3), 491–500.

Greenman, C. D. *et al.* (2010). PICNIC: an algorithm to predict absolute allelic copy number variation with microarray cancer data. *Biostatistics*, **11**(1), 164–175.

Grossman, R. L. *et al.* (2016). Toward a Shared Vision for Cancer Genomic Data. *The New England journal of medicine*, **375**(12), 1109–1112.

Hofmann, M. H. *et al.* (2021). BI-3406, a Potent and Selective SOS1–KRAS Interaction Inhibitor, Is Effective in KRAS-Driven Cancers through Combined MEK Inhibition. *Cancer Discovery*, **11**(1), 142.

Li, H. *et al.* (2009). The Sequence Alignment/Map format and SAMtools. *Bioinformatics*, **25**(16), 2078–2079.

Mayrhofer, M. *et al.* (2016). Rawcopy: Improved copy number analysis with Affymetrix arrays. *Scientific reports*, **6**, 36158–36158.

McDonald, E. R. *et al.* (2017). Project DRIVE: A Compendium of Cancer Dependencies and Synthetic Lethal Relationships Uncovered by Large-Scale, Deep RNAi Screening. *Cell*, **170**(3), 577–592.e10.

Meyers, R. M. *et al.* (2017). Computational correction of copy number effect improves specificity of CRISPR–Cas9 essentiality screens in cancer cells. *Nature Genetics*, **49**(12), 1779–1784.

Munzner, T. (2014). *Visualization Analysis and Design*. CRC Press, Taylor & Francis Group.

Nassar, A. H. *et al.* (2021). Distribution of KRASG12C Somatic Mutations across Race, Sex, and Cancer Type. *New England Journal of Medicine*, **384**(2), 185–187.

Rogers, J. *et al.* (2019). Composer — Visual Cohort Analysis of Patient Outcomes. *Applied Clinical Informatics*, **10**(02), 278–285.

Satyanarayan, A. *et al.* (2017). Vega-Lite: A Grammar of Interactive Graphics. *IEEE Transactions on Visualization and Computer Graphics*, **23**(1), 341–350.

Zhang, Z. *et al.* (2015). Iterative cohort analysis and exploration. *Information Visualization*, **14**, 289–307.
